# Programmable Assembly and Steering of Microbubble Droplets using Ultrasound

**DOI:** 10.1101/2025.01.08.631861

**Authors:** Alexia Del Campo Fonseca, Khemraj Gautam Kshetri, Dimitar Boev, Nathan Irniger, Nitesh Nama, Daniel Ahmed

## Abstract

Gas-filled microbubbles have been extensively used as contrast agents for ultrasound therapies and have recently been explored as microrobots. When exposed to an intense acoustic wave, microbubbles oscillate and scatter the sound field, leading to assembly and manipulation behaviors. Although traditional microbubbles demonstrated promising potential for manipulation in physiological environments, we recognized the need for improved assembly selectivity and acoustically directed propulsion. For this, we developed a new microbubble design using microfluidics. We engineered microbubbles with an outer oil droplet encapsulating a gas core, allowing the gas to move freely within the oil. When the gas reaches the droplet’s periphery, the oil attenuates the scattered wave, ensuring that scattering or acoustic amplification is concentrated near the droplet’s edge. This approach enabled us to control the organized assembly of microbubbles and direct their acoustic navigation. We believe this approach will open new possibilities for using gas-filled microbubbles as safer drug carriers or for micro-interventions in biomedical settings.

## Introduction

The manipulation of microbubbles through acoustic means has garnered significant attention, particularly in the realm of biomedical research (*1*). Commercial microbubbles are micron sized gas filled lipid spheres that are meant to navigate the circulatory system and provide strong scattering of the ultrasound coming from outside of the body. Currently, they are widely used in clinical settings as ultrasound imaging contrast agents (*2*–*5*). Additionally, they have also been investigated as drug delivery microrobotic platforms (*1, 6*–*10*), to aid manipulation of microstructures such as polymeric microrobots (*11*–*18*) or to induce streaming for applications such as biofilm removal (*19*). Overall, the small size, low toxicity, and high responsiveness to sound make microbubbles a promising approach for non-invasive and high-precision therapies (*5, 20, 21*). However, while their high acoustic response is their main strength that promises their external manipulation even inside deep tissue regions, there is currently a lack of precise control over this process. When exposed to an acoustic field, microbubbles scatter sound waves in all directions that, in turn, interact with their three-dimensional (3D) environment in a largely uncontrolled and indiscriminate manner. This results in disordered microbubble assemblies and unpredictable propulsion behaviors (*22*– *25*). Efforts to fully control the cascade of events that occur when microbubbles are within a sound field will significantly enhance microrobotic approaches facilitating their application and integration into physiological environments.

The current generation of microbubbles typically utilizes a core hydrophobic gas (i.e. SF6, perfluorocarbons, etc) to improve gas stability in the surrounding hydrophilic media (*3, 23, 26*). This core is encased in a surrounding layer of molecules (i.e., phospholipids, polymers, proteins …) designed to be flexible and as thin as possible, thereby minimizing acoustic damping and promoting the scattering of sound(*26, 27*). When subjected to an acoustic field, the gas within microbubbles readily compresses and expands, resulting in the amplified scattering of sound waves. This scattering not only amplifies the primary radiation forces on the microbubbles, driving them in the direction of wave propagation, but also generates secondary pressure-based forces between neighboring microbubbles, leading to their aggregation into clusters (*22, 24, 28*–*30*). Additionally, the transmission of scattered waves through the fluid creates acoustic streaming patterns, capable of producing forces strong enough to propel even polymeric structures (*13, 31*–*33*). However, current microbubble-based propulsion lacks directionality and often relies on additional actuation modalities, such as magnetic fields, for microrobotic steering (*11, 12, 34*). Developing a purely acoustic steering and propulsion mechanism would greatly enhance their application as microrobots. Moreover, the aggregation process typically occurs rapidly, with microbubbles continuously shifting positions due to dynamic self-attraction. Achieving controlled assembly could unlock the potential to create customizable microbubble swarms, where inner microbubbles could carry specific drugs while outer layers transport different molecules.

We present an alternative microbubble design that deviates from conventional approaches by fabricating gas bubbles that hover inside an oil droplet. These microbubbles allow the gas to move freely within the oil, directing acoustic energy toward the gas location. This design promotes the formation of ordered configurations, with the microbubbles quickly organizing into highly structured arrangements, such as doublets, triplets, or crystalline-like formations, through self-assembly. Additionally, changes of direction of the traveling acoustic wave lead to changes in direction of the inner gas, determining the subsequent propulsion directionality. This effect allows for precise steering of the microbubbles’ movement. The development of this new approach has the potential to usher in a new era of highly controlled microbubbles.

## Results

### Experimental set-up

This study comprises two main tasks: high throughput fabrication of oil-gas microbubbles and its investigation under acoustic influence. For microbubble fabrication, we designed a microfluidic device that guides the flow of three different fluids coming from three separate inlets on the device (*35*). The channels converge at a junction where a three-phase emulsion of gas-oil-water forms (see **Fig. 1**). We transferred the 2D design on to a silicon wafer through a process of photolithography, see Materials and Methods. We used the resultant silicon wafer as a mold for a subsequent soft lithography process. A mixture of polydimethylsiloxane (PDMS) Sylgard base composite and Sylgard curing agent in a 10:1 ratio was poured over the wafer, degassed and cured in the oven at 85°. After demolding, the PDMS slab was punched 4 times, 3 holes for inlets and 1 hole for the outlet. Finally, the PDMS was attached via plasma treatment to a glass slide. **Fig. 1a, d** shows a top-view of the chip during fabrication. The output section of the microfluidic device, where the produced microbubbles flow, was treated with polyvinyl alcohol (PVA) to render the PDMS hydrophilic and prevent oil attachment to the walls (see full procedure for PVA coating in Materials and Methods).

**Fig. 1.**
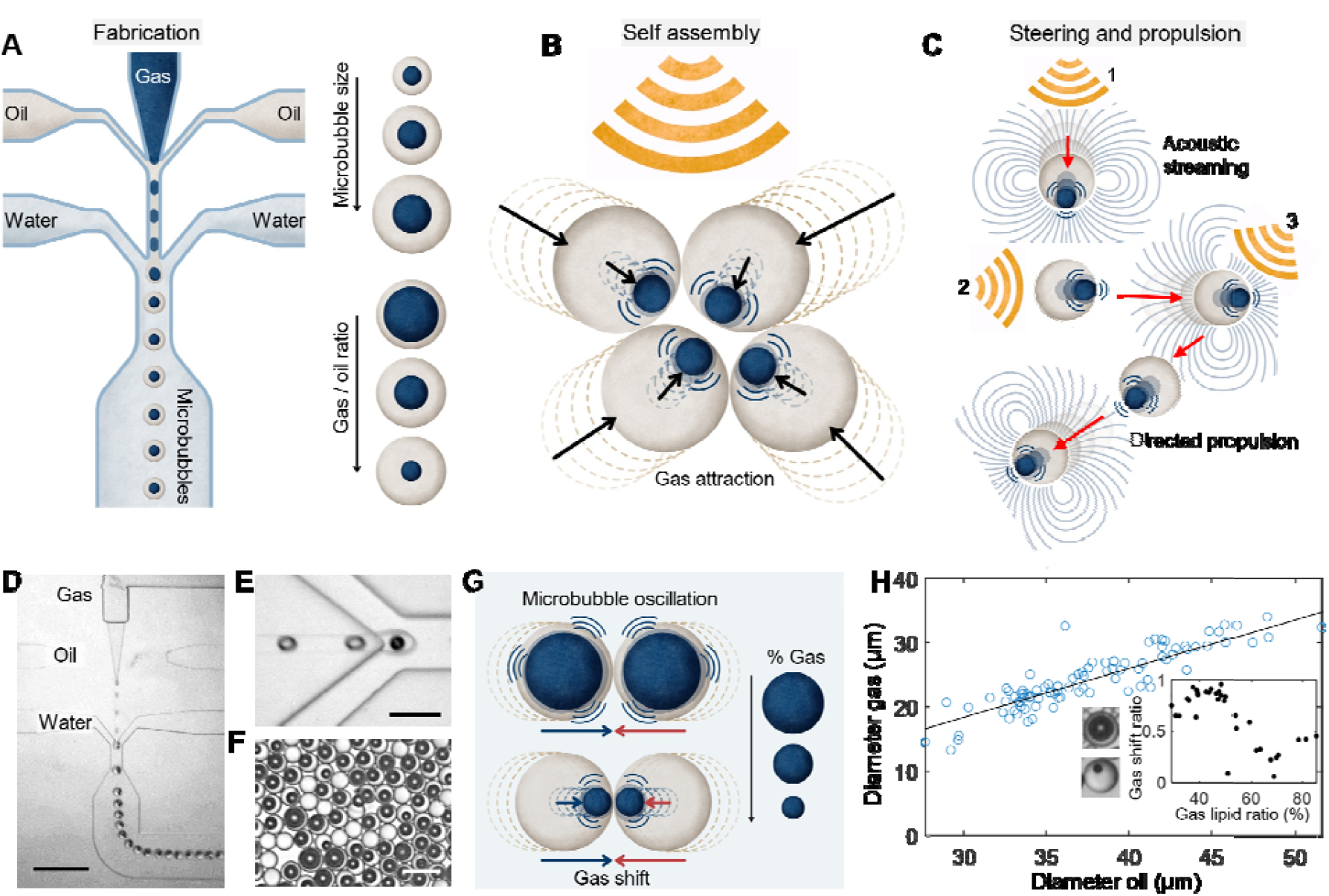
Acoustic droplets fabrication and response to ultrasounds. **(A)** Microfluidics fabrication of acoustic droplets. A double emulsion is generated featuring water, oil and gas. The control of flow rates of each fluid allows for control of the microbubble size and oil vs gas ratio, as shown on the right of the schematic. **(B)** Acoustic droplets inside an ultrasound field attract towards each other, forming ordered assemblies. **(C)** Ultrasound coming from different directions allow for the steering of droplets in space. Additionally the appearance of acoustic streaming, leads to self-propulsion of this droplets inside a fluid. **(D)** Microscope image snapshot of the microfluidic fabrication process, and the formed acoustic droplets. Scale bar is 100 µm. **(E)** Zoomed microscope image of the right moment where a double emulsion is being formed. Scale bar is 30 µm. **(F)** Population of formed acoustic droplets imaged with optical microscope. Scale bar is 50 µm. **(G)** Schematic showing the role of gas vs oil ratio. Lower percentage of gas means higher motion of the gas inside the droplet. **(H)** Plot representing the correlation between gas size and oil size. The line is a fitting straight line to show the increasing tendency of the plot. Inset: Correlation between gas movement (y axis) and gas-lipid ratio. Lower gas-lipid ratio turns into higher gas movement.

For emulsion generation, three distinct fluid flows were introduced into the microfluidic device through separate inlets (*35*) (**Fig. 1a, e**). The first flow (water) consisted of water mixed with P188 Poloxamer polymer and glycerol (see **Table 1**). The second flow (oil) contained octanol-1. These two flows were guided into the microfluidic channels using Tygon tubing connected to programmable syringe pumps. The third flow comprised gas SF6 with 3% purity, directly sourced from a gas cylinder and regulated by two pressure regulators before entering the system via Tygon tubing. The fabrication setup was mounted on an optical microscope and monitored using high-sensitivity cameras. The fabricated microbubbles were transferred to a single straight-channel microfluidic device. The PDMS channel diameter was 400 µm and the height was 30 µm. This setup enabled the examination of microbubble behavior under acoustic stimulation. To facilitate this, we affixed a piezoelectric transducer to the side of the PDMS using a two-component epoxy adhesive. The transducer was connected to a function generator, and the set-up was installed on an inverted microscope. The behavior of the microbubbles was captured using high-sensitivity cameras.

**Table.1.** Emulsion mixture recipes.

| **Outer aqueous solution (OA)** |  |
| --- | --- |
| **Glycerol** | 1.5 ml |
| **Water** | 3.5 ml |
| **Poloxamer 188 (P188)** | 5 ml |
| **Lipid Octanol solution (LO)** |  |
| **Lipid working solution (DOPC)** | 0.1 ml |
| **1-octanol** | 4.9 ml |

### Fabrication of Gas-Oil Droplets using microfluidics

Traditionally, microbubbles contain a gas core which is coated by a single layer of polymer or lipid molecules, mainly to promote gas stabilization. During this study, we explored an unconventional microbubble design. We explore the fabrication of emulsions where the gas appears in the form of a bubble that hovers inside an oil droplet. Note that we use the term ‘bubble’ to refer to gas components and ‘droplet’ to refer to fluid components. The oil droplets were formed by emulsification inside an aqueous fluid, while gas was simply introduced into the oil medium **Fig. 1e, f**. This configuration introduces a new level of freedom during the fabrication of microbubbles. It allows for precise control over the size of the gas bubble while ensuring the overall droplet size remains undisturbed. The size of the oil droplet is determined by the flow rates used during microfluidic fabrication. By using flows of 10 to 12 µl/min for the water solution and 1 to 2 µl/min for the oil solution, we fabricated droplets between 20 µm and 50 µm. On the other hand, the amount of encapsulated gas is controlled by regulating the pressure coming from the gas cylinder. We used pressures between 0.2 and 1 bars and we obtained encapsulated bubbles that range between 10 µm and 40 µm. The size distribution that we used during experiments is represented in **Fig. S1**. While we kept the size of the oil droplet relatively constant during emulsion fabrication, the size of the gas bubble encapsulated inside it varied on our demand (see **Fig. 1a**).

During microbubble fabrication, we explored the effect of gas content on microbubble acoustic response. The mathematical expressions describing acoustic radiation forces on bubbles indicate that larger bubbles experience larger forces, suggesting that higher gas content should lead to stronger radiation forces and better acoustic responsiveness (*25, 36*). However, we discovered that the relative size of the gas to oil ratio is also crucial in determining acoustic behavior. We measured the diameter of gas bubble with respect to the diameter of the oil for each droplet. Here, we observed that with high gas/oil ratio, these emulsions behave as standard microbubbles. They assembled into clusters and propelled, as seen in literature (*25, 36*). However, as the percentage of gas is decreased, the droplet creates room for the gas bubble to undergo relative motion within the oil. The combination of buoyancy force and the tendency towards minimum free energy, drives the gas bubble to rest at the center of the droplet, under static conditions, see **Fig. 1**. When ultrasounds are introduced in the system, the gas bubble inside the oil, guided by acoustic radiation forces, shifts its place from the center to one side of the droplet. We measured the displacement of the gas bubble from the center of the droplet, under acoustic excitation, see **Fig. 1h**. In **Fig. 1h**, 0 represents the center of the droplet and 1 indicates the scenario where the bubble comes into contact with the edge of the droplet. We observed that in the range from 60% to 50% of gas-oil ratio, there is a transition between no gas displacement to total gas displacement, see **Fig. 1h**. The bubble displacement triggers a cascade of assembly and propulsion events which differ from traditional microbubbles, as illustrated in **Fig. 1c, d**. These events will be further discussed in the following sections.

### Selective assembly of microbubble droplets

Microbubbles and droplets subjected to an acoustic field are known to experience primary acoustic radiation force in the direction of a traveling acoustic wave. In our system, owing to the relatively larger acoustic impedance mismatch between gas and oil, the gas bubble experiences a larger primary radiation force. As such, the inner gas-oil interface moves faster than the outer oil-water interface. Consequently, for an isolated gas-oil droplet, the gas moves in the direction of acoustic wave propagation to occupy an off-centered position within the oil droplet. In the presence of neighboring droplets, the gas-oil droplet also experiences an attractive secondary Bjerknes force in addition to the primary acoustic radiation force. This secondary Bjerknes force scales inversely with the second power of the inter-bubble distance. If the droplets are sufficiently close, the secondary Bjerknes force is significantly larger than the primary radiation force, resulting in their motion towards each other and the self-assembly of gas-oil droplets(*37*), **Fig. 2a**. To delve into this matter, we begin by examining the system’s initial response to the acoustic wave, focusing on a group of five bubbles. Their sizes and movements over time are depicted in **Fig. S2**; note that the colors in **Fig. S2** correspond to those in **Fig. 2**, offering additional and more detailed analysis.

**Fig. 2.**
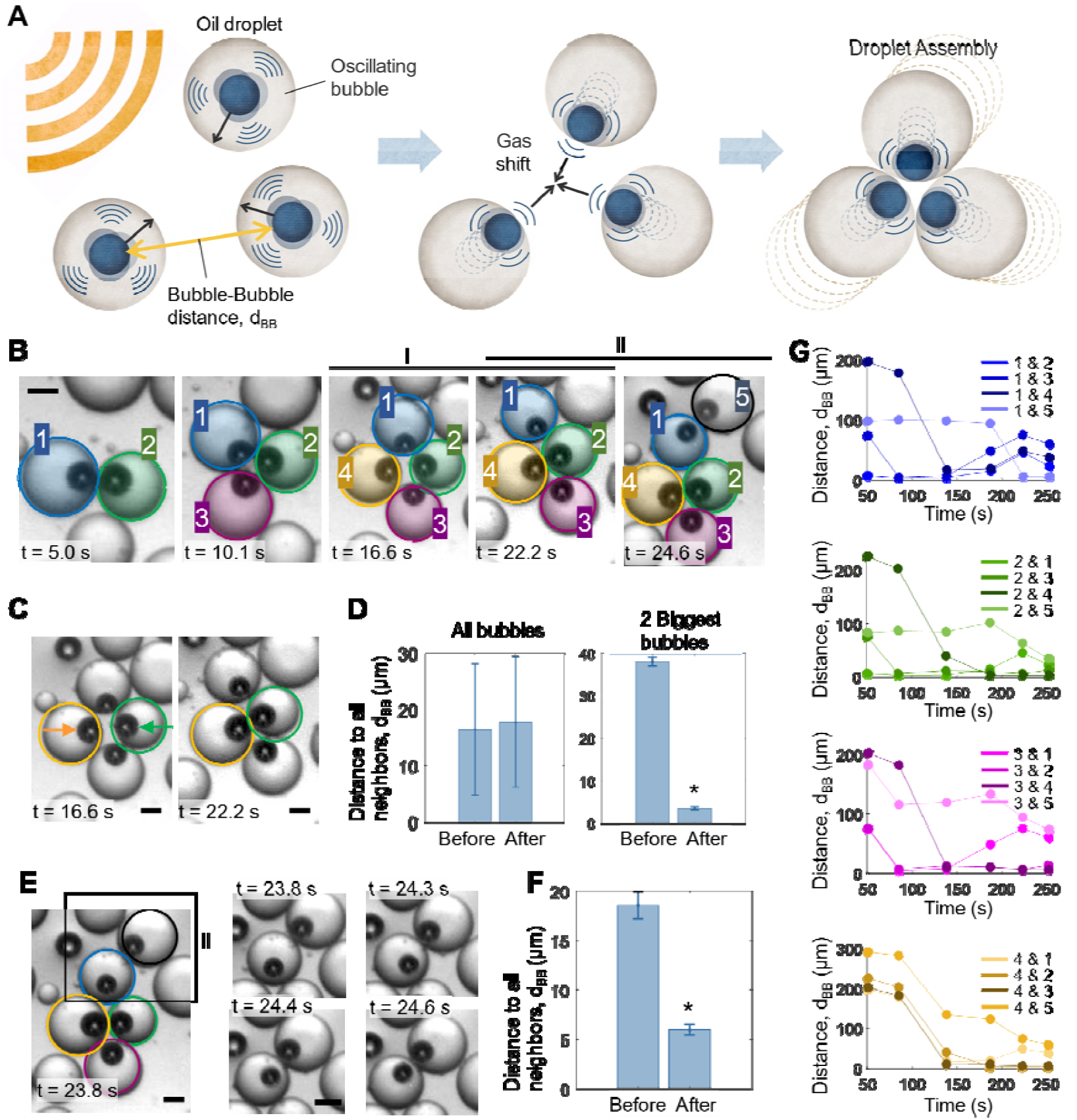
Microbubble droplets assembly into ordered clusters. **(A)** Schematic of the assembly process. Inner gas bubbles are attracted to neighboring bubbles. After the initial bubble motion, the droplets are also dragged to form organized assemblies. **(B)** Experimental images of droplet assemblies with 2, 3, 4 and 5 bubbles. The various assemblies appear over time, while the droplets are under the influenced of an ultrasound field of 242 kHz and 20 V_PP_. The colors used correspond to the colors shown in ‘G’. Scale bar is 10 µm. **(C)** These experimental images show the strong influence of bigger bubbles in the final assembly configuration. Bigger bubbles (marked in orange and green) will attract to each other strongly, producing a rearrangement of the cluster, and determining the final assembly shape. **(D)** First plot (left) measures the average of bubble-bubble distance, between all the bubbles of the system, before and after the cluster shape rearrangement. Second plot (right) measures the average of bubble-bubble distance, between only the two biggest bubbles of the system, before and after the cluster shape rearrangement. **(E)** These experimental images show the influence of an incoming bubble to an already established assembly. Images on the right show the sequence of events over time, where a bubble inside the cluster, reorients to face the new bubble. This event changes the cluster configuration. **(F)** This plot measures the average of bubble-bubble distance, between all the bubbles of the system, before and after the cluster rearrangement. **(G)** These plots measure bubble-bubble distance between all bubble pairs in the system, over time. Each color represents one bubble (also identified with numbers from 1 to 5).

In these experiments, we observed that upon acoustic activation, the encapsulated gas bubble moved away from the center of the droplet towards the edges, specifically in the direction of the nearest neighboring encapsulated gas bubbles, see **Fig. 2a**. Additionally, we observed that the gas within the droplet can also displace the entire droplet in the same direction as the initial gas movement, **Fig. 2a**. Consequently, in situations involving two neighboring bubbles, both droplets move toward each other driven by the acoustic forces acting between their encapsulated gas. This phenomenon results in the self-assembly of microbubble droplets into clusters, **Fig. 2b**. We measured the speeds of droplets on their movement towards each other in **Fig. S3**. Note that our analysis was conducted within a microfluidic device of 30µm in height, thus it only allowed for a two-dimensional layer of bubbles; this helped with the visualization of microbubble assembly in groups, **Fig. 2b**. Subsequently, when looking to a group of microbubble droplets, upon acoustic activation every gas bubble either converged towards the cluster’s center or aligned itself towards it. This behavior, combined with the steric effects from the surrounding uniformly sized oil droplets, leads to the formation of highly ordered arrangements, **Fig. 2a, b**.

It’s important to note that natural self-assemblies are normally guided by the principle of minimizing free energy. Similarly, our microbubble droplets consistently moved towards positions of least distance from their neighboring bubbles, effectively diminishing the system’s overall free potential energy. We examined combinations of 2, 3, 4, and 5 bubbles, tracking their evolution in terms of bubble-to-bubble distances. We show an example in **Fig. 2c, d**, where 4 bubbles rearrange such that the bigger bubbles are in close contact with each other, keeping the other two as close as possible. Notably, larger bubbles exert stronger attraction forces, leading to reduced distances and lower energy levels, **Fig. 2d**. In some instances, larger bubbles approaching an existing cluster can induce rearrangements within the cluster, causing the closer bubble to re-orient outward towards the larger incoming bubble, see **Fig. 2e, f**. Subsequently, we conducted an analysis of the nearest distance to a neighboring bubble for each droplet within the system, as depicted in **Fig. 2g**, over time. For this characterization, we selected four bubbles corresponding to the four graphs in **Fig. 2g** and tracked their distance to all other bubbles in the system over time. Across all cases, we observed a trend towards minimizing the distance between bubbles, with larger bubbles, see **Fig. 2g**, playing a prominent role in energy minimization.

Overall, despite experiencing identical acoustic excitation, bubbles can dynamically alter their positions, guided by their closest and most efficient interactions within the surrounding environment. The tendency to cluster, driven by the principle of minimizing free energy, along with the attenuation of the scattered sound by the surrounding oil droplet, the uniform size of the droplets, and the steric effects imposed by the oil, collectively results in highly ordered assemblies.

### Steering mechanism of microbubble droplets

Microbubbles are known not only to self-assemble but also to navigate through space, driven by primary radiation forces exerted by incident acoustic waves in the direction of wave propagation. Specifically, when the microbubble droplets immersed in a surrounding liquid are subjected to an acoustic field, both the gas-oil and the oil-water interface oscillate, leading to a net time-averaged acoustic radiation force and the resulting propulsion of gas-oil droplets. The net force experienced by a gas-oil droplet can, in general, include contributions from the primary acoustic radiation force (arising due to scattering of applied acoustic field), the secondary Bjerknes force (arising due to interactions with neighboring gas-oil droplets), and the microstreaming created due to viscous dissipations of acoustic waves near the oscillating interfaces (*37*). Combined, the total hydrodynamic force experienced by the gas-oil droplet is given by 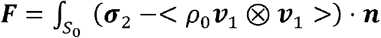 with 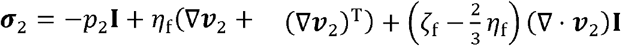, where *p*_2_ indicates the time-averaged hydrostatic pressure, the terms containing *ν*_2_ arise from the microstreaming, and the term containing *ν*_1_ indicates the nonlinear interactions of the acoustic wave with itself. Our studies have demonstrated that by strategically positioning transducers at various angles relative to the microbubbles, we can control their movement in 3D directions and guide them through intricate vascular networks, such as those in the brain(*36*). Building on this knowledge, we have utilized radiation forces to induce steering in microbubble droplets. When we activate the acoustic field, the initial movement of the gas bubble within the oil droplet serves as a calibration for the direction of movement, see **Fig. 3a**. By positioning a transducer on the left, the radiation force causes the gas bubble to move towards the right edge of the droplet, **Fig. 3a**. Once this alignment is achieved, the droplet begins to self-propel in a directed movement towards the right, **Fig. 3a**.

**Fig. 3.**
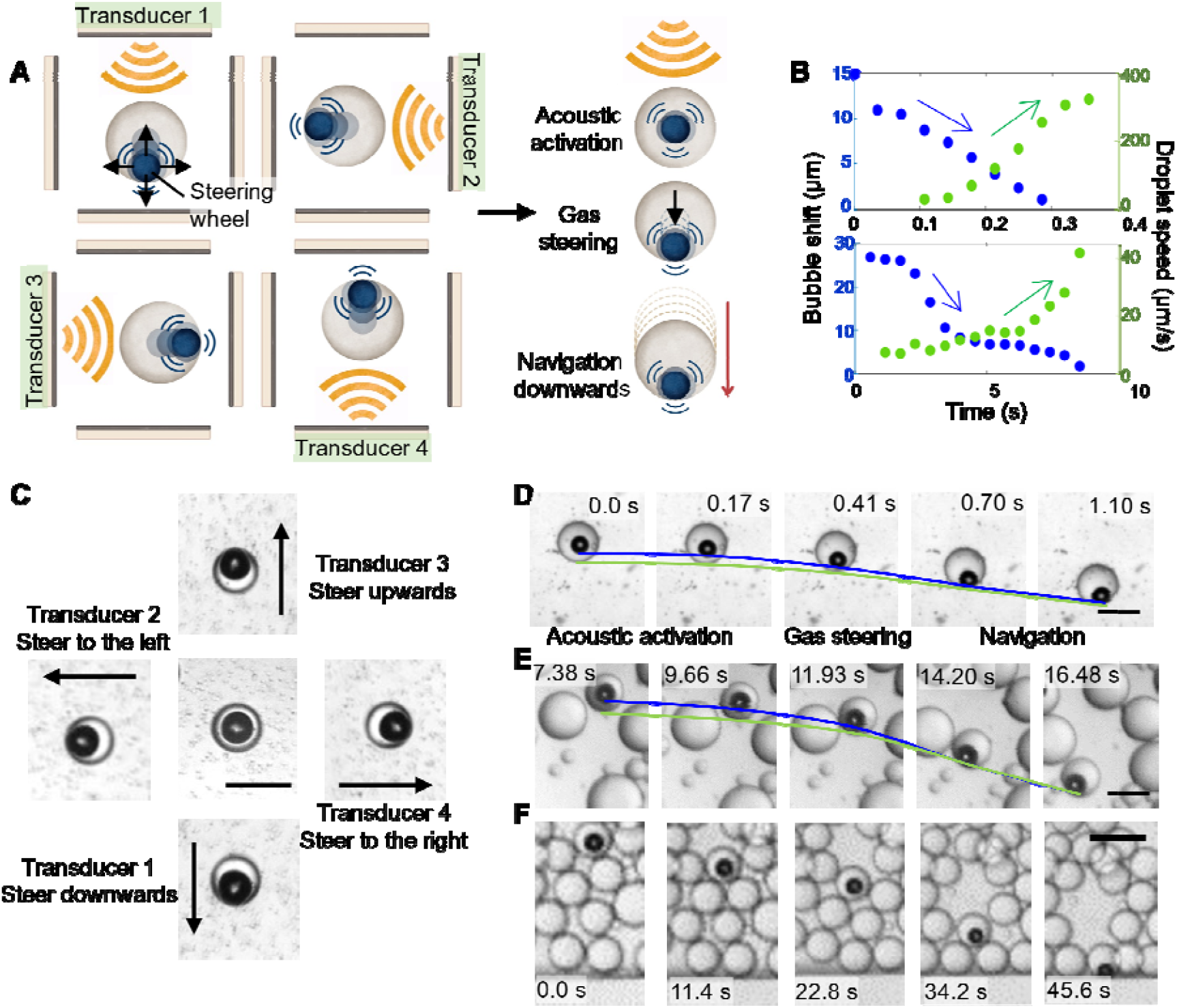
Acoustic droplet steering using primary radiation force. **(A)** Schematic representation of the steering principle. The gas shifts from the center of the droplet in the direction of wave propagation, thereby determining the direction of subsequent propulsion. Transducers positioned at various orientations enable the steering of microbubble droplets. **(B)** Correlation over time between inner gas motion to the edge of the droplet (bubble shift, in blue) versus speed of droplet manipulation (in green). **(C)** Schematic of the steering principle shown with real images of the microbubble droplets. Scale bar is 20 µm. **(D, E)** Sequence of microscope images in which we see a droplet under ultrasounds over time. In blue line we track the movement of the inner gas and in green line the movement of the droplet’s edge. Scale bar is 30 µm. **(F)** Sequence of microscope images in which we see a droplet under acoustic excitation, moving towards a boundary. Scale bar is 50 µm.

To characterize this steering principle, we placed four piezoelectric transducers around the experimental PDMS chamber in an orthogonal arrangement, allowing for actuation in the four primary directions: up, down, left, and right, **Fig. 3a**. We fabricated microbubbles and then excited them sequentially with each piezoelectric transducer individually. In each case, we observed that the gas initially relocated towards the direction of wave propagation (up, down, left, or right, respectively), while the oil droplet remained static. Subsequently, if the acoustic field was sustained, the droplet began to gain speed and moved in the direction of the initial gas steering, see **Movie S1 and Movie S2**. We characterized this event by measuring the position of the gas bubble inside the droplet over time and simultaneously measuring the speed of the entire droplet in the direction of wave propagation (see **Fig. 3b**). In **Fig. 3b**, the value y_max_, represents the center of the droplet, while y = 0, indicates the point where the bubble touches the edge of the droplet, as observed through the optical microscope. In the initial moments of gas motion, we observed that the droplet’s speed was close to zero. However, as the gas bubble moved significantly towards the side, the droplet began to accelerate. **Fig. 3b** illustrates this phenomenon with two examples, each represented in separate plots. Additionally, **Fig. 3d, e** presents the experimental image sequence. In these images, the movement of the gas bubble is marked with a blue line, while the movement of the oil droplet is marked with a green line. Over time, these lines converge as the gas bubble approaches the edge of the oil droplet. We also demonstrate this phenomenon in the presence of a boundary, **Fig. 3f and Fig. S4**. When a wall is nearby, the microbubble is attracted to it due to secondary Bjerknes forces (*30*).

### Propulsion of microbubble droplets

In addition to steering and directed motion, these microbubbles also exhibit propulsion. As extensively studied in the literature, microbubbles oscillate and strongly scatter sound when excited by acoustic waves, generating surrounding streaming flows. Acoustic streaming refers to the steady fluid flow induced by the inertial rectification of acoustic transmission through a viscous fluid.

In our oil-gas droplet system, the dissipation of acoustic waves within the boundary layers of both the gas-oil and oil-water interfaces results in a time-averaged streaming flow, both within the oil and surrounding liquid. **Fig. 4a** and **Fig. S5** illustrate the initial motion of microbubble within the droplet resulting from the primary acoustic radiation force, and the accompanying acoustic streaming patterns around the gas-oil droplet. To visualize the streaming patterns around the gas-oil droplet, we introduce tracer particles of size 1 µm and track their trajectories. Our experimental observations in **Fig. 4b** indicate a monopole streaming pattern characterized by two streaming rolls around the gas-oil droplet. These patterns indicate that the microbubble interface oscillates primarily in a combined volumetric and translational mode and are in excellent agreement with our numerical predictions shown in **Fig. 4c**, (*38*).

**Fig 4.**
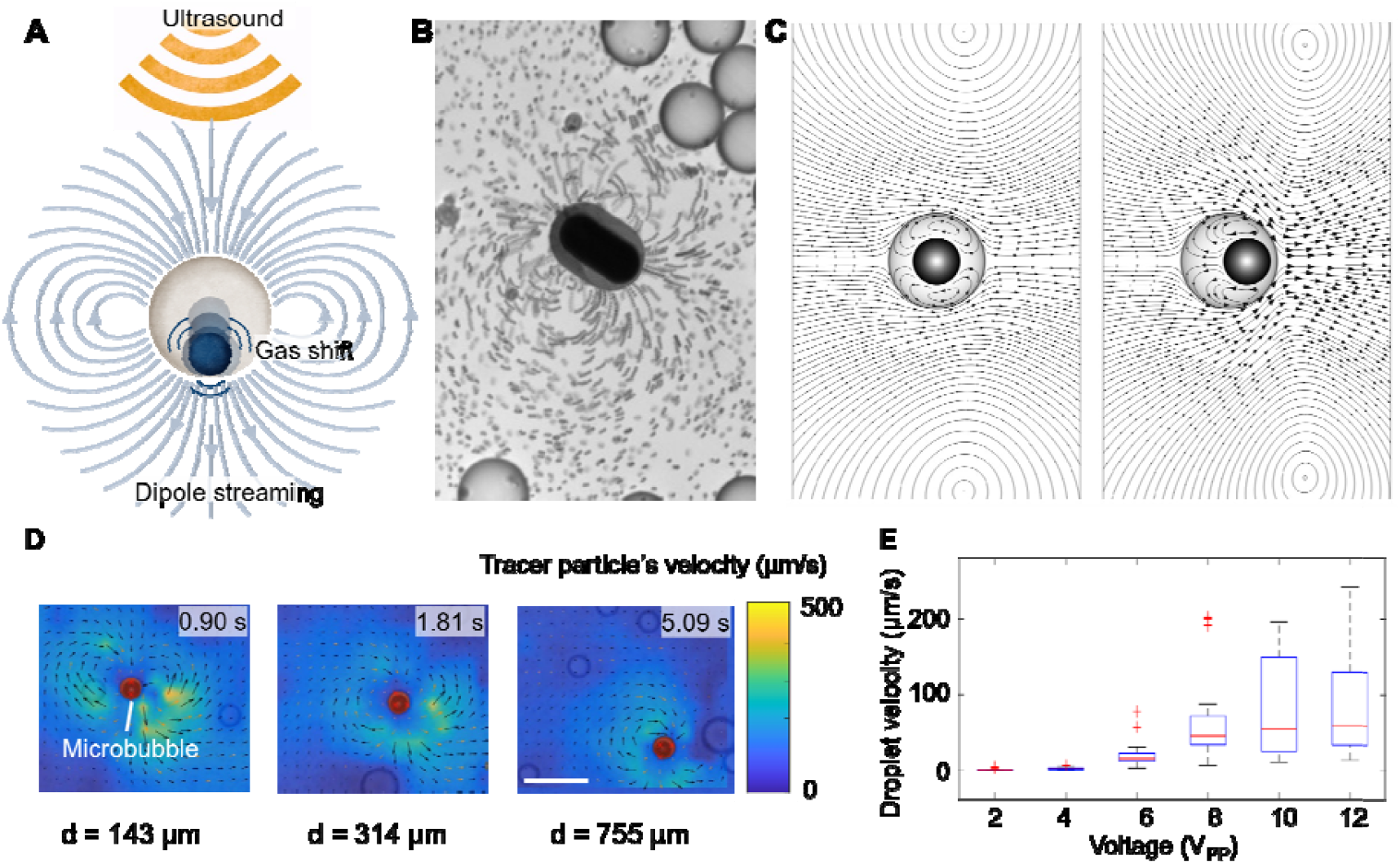
The role of acoustic streaming for microbubble droplet propulsion. **(A)** Schematic representation of the acoustic streaming generated around microbubble droplets, where the inner gas occupies an off-centered position. A dipole-shaped streaming is generated, and the droplet moves in the direction of wave propagation. **(B)** Superposition of microscope images over time, showing the generation of dipole shape streaming around a microbubble droplet. 1 µm tracer particles have been located in the surrounding fluid, in order to visualize the streaming flows. **(C)** Streamlines of mean velocity around a single microbubble droplet obtained with numerical simulation. The left plot shows the streaming field around the microbubble droplet when bubble is placed at the center of the droplet while right plot shows the streaming field when bubble occupies off-centered position within the droplet. The size of the arrow in these figures is indicative of the respective streaming velocity magnitude and reveals significantly higher streaming velocities for the off-centered bubble placement. **(D)** Image sequence showing the PIV analysis performed on the tracer particles that have been placed in the fluid surrounding the droplet. Each image is taken at an increasing distance from the transducer (from left to right). A deeper analysis is shown in Fig. S5. **(E)** Plot representing the effect of voltage in the system. Higher voltage applied to the piezoelectric transducer, results in higher droplet velocities.

To further assess the effect of gas motion within the droplet on the resulting microstreaming, we also performed numerical simulation by placing the bubble in an off-centered position. Comparing our results against the case with symmetric placement of bubble in **Fig. 4c**, we observed that the off-centered motion of gas within the droplet leads to the shifting of microstreaming vortices in the direction of gas motion. The size of the arrow in these figures is indicative of the respective streaming velocity magnitude and the maximum streaming speed is observed to increase as the bubble moves away from the center of the droplet.

Further, we observed that as the microbubble moves further from the transducer, the streaming speeds decrease in intensity, see **Fig. 4d, Fig. S5** and **Movie S3**. We additionally characterized this phenomenon at different voltages applied to the piezoelectric transducer, see **Fig. 4d**. Increased voltage results in greater vibration of the piezoelectric material, producing sound waves of higher amplitude. Consequently, higher amplitude corresponds to stronger radiation forces and stronger microbubble oscillations. Our findings confirm that microbubbles move faster with higher activation voltages, see **Fig. 4b**. Overall, we observed that the generation of acoustic streaming around microbubble droplets leads to a purely acoustic propulsion mechanism. Notably, when combined with the previously discussed steering principle, this effect results in a directed acoustic propulsion approach, see **Movie S4**.

## Discussion

We have fabricated gas-filled oil droplets that respond efficiently to acoustic forces in a controlled fashion. These microbubbles consist of an outer oil droplet surrounding an inner hydrophobic gas bubble, both suspended in a water background. This design allows the encapsulated gas to move freely within the oil droplet, which is the key feature enabling controlled self-assembly and directed propulsion.

We have demonstrated that the oil-to-gas ratio is crucial in determining both the stability of the bubble over time and the magnitude of the acoustic forces it generates. Lower gas-to-oil ratios allow for the free motion of gas within the oil, forming the desired droplets for our study. However, if the gas-to-oil ratio is too low, the gas will collapse, and the droplet will not respond to the acoustic field, see **Fig. S4**. Therefore, precise control over the amount of gas and oil during microbubble fabrication is essential. To achieve this control, we utilized microfluidics. This technology, previously shown to be effective for forming emulsions, proves capable of creating gas-filled oil droplets. Microfluidics allows for precise adjustment of oil and gas components by tuning the inlet pressure of each fluid. It also facilitates the incorporation of additional components into either the coating or the gas content of the droplet.

Once the microbubbles were fabricated, we demonstrated their assembly as an organized group. When the acoustic field was activated, our specially designed droplets exhibited gas relocation in the direction of the nearest neighboring droplet. Subsequently, assembling into a cluster. We confirmed that the droplets move towards configurations of minimum free energy, which corresponds to the minimum distance between neighboring microbubbles. This phenomenon, combined with the steric effect of the oil droplets and the attenuated scattered waves on the outer part of the droplet, resulted in the formation of highly ordered clusters.

Additionally, we demonstrated the steering and navigation capabilities of these droplets during their movement in space. The free motion of the gas within the droplet allows for redirection of the droplet’s movement. By positioning various piezoelectric transducers, we can relocate the gas to different edges of the droplet, thereby controlling the desired propulsion direction. We also observed that droplet motion was driven partly by the primary radiation force exerted by the piezoelectric transducer and partly by the streaming propulsion forces generated during microbubble oscillations. We characterized the streaming pattern of these droplets as having a dipole shape, which contributes to maintaining a determined direction during motion. Finally, we confirmed that droplet speeds are controllable by tuning the voltage supplied to the function generator.

These new microbubbles still require optimization. Their stability over time needs further improvement, and alternative approaches such as polymer coatings should be explored. Nevertheless, these microbubbles show significant potential to overcome the limitations of commercially available microbubbles by including high fluid volumes around the droplets, which can be used for drug loading, Also, protecting the external environment from the acoustic effects of vibrating objects. This is achieved by covering the vibrating objects (in this case, microbubbles) with a thick oil layer, which partially dampens the reflected acoustic waves. Additionally, we demonstrated a technique that imposes order on the self-assembly process of microbubbles. And we introduced a purely acoustic propulsion approach, which includes acoustic steering, eliminating the need for other actuation mechanisms, such as magnetics. This platform opens up new possibilities in the microbubble field, allowing for the exploration of various designs that can alter their acoustic response and even facilitate their control, ultimately enabling their use in physiological conditions.

## Materials and Methods

### Microfluidic device fabrication

The microfluidic devices were designed using Layout Editor, a 2D computer-aided design software. The 2D microchannel patterns were transferred onto a photomask via laser writing, which was subsequently used for photolithography. In this process, a silicon wafer was coated with SU8-3050, aligned with the photomask, and exposed to UV light. After exposure, the wafer was developed to create a positive mold of the desired microchannel pattern. The finished wafer was then vapor-coated with silane. For microfluidic device fabrication, PDMS Sylgard base elastomer was mixed with Sylgard curing agent at a 10:1 ratio and poured onto the wafer. The mixture was placed in a vacuum chamber to remove trapped gas bubbles by maintaining the vacuum for approximately 20 minutes. The uncured PDMS was then cured in an oven at 85°C for 90 minutes. After curing, the PDMS was removed from the mold, cut into individual devices, and holes were punched with a 0.75 mm diameter puncher to serve as inlets and outlets. The devices underwent a cleaning process to remove any contamination. Adhesive tape was initially applied to both sides to eliminate dust and particles. Subsequently, the devices were immersed in isopropyl alcohol (IPA) and placed in an ultrasonic bath for 10 minutes. After cleaning, the devices were dried with pressurized nitrogen and attached to glass slides via plasma bonding. Both the glass and the PDMS inner surfaces were treated simultaneously with a plasma beam for 30 seconds each and then bonded together. To solidify the attachment, the bonded devices were heat-treated in an oven at 85°C for an additional 90 minutes.

### PVA coating

PVA powder from Sigma-Aldrich was mixed with deionized water at a 2.5‰ wt./vol ratio, stirred at 500 rpm, and heated to 80°C until dissolved. The PVA solution was introduced into the device through the outlet, while gas was simultaneously injected from the gas inlet to prevent PVA overflow into other channels. This flow was maintained for 3-5 minutes. Excess PVA was removed by increasing the airflow rate, and the devices were heated at 120°C for 15 minutes to eliminate residual moisture.

### Emulsion mixtures

The device featured three inlets (from top to bottom): The first inlet was connected to an external aqueous solution (OA). The second inlet was linked to a lipid-containing 1-octanol solution (LO). The third inlet was for gas (in our case, SF6). At the intersection, the three phases were combined to generate a double emulsion, subsequently matured to form a gas-filled liposome. DOPC obtained from Avanti Lipids served as the lipid working solution and required storage at -20°C. Poloxamer 188 (P188), Glycerol and octanol were procured from Sigma-Aldrich. SF6 was purchased as a 10L gas cylinder with working pressure between 0.3 and 3 bar, from Linde. The composition of the phases is detailed in the table provided below.

### Acoustic activation

To investigate the acoustic response of microbubbles, we fabricated a dedicated microfluidic device. This device featured a single channel measuring 400um in diameter and 30um in height. The inlet of this device was linked to the outlet of the fabrication device through a custom designed metallic connector, bent to fit into both openings. This setup ensured that the fabricated microbubbles could flow directly into the new device for observation and subsequent acoustic activation. For acoustic activation, a piezoelectric transducer was employed. We utilized a transducer sourced from Stemminc, comprising a ceramic piezo plate measuring 7×8×0.2 mm. The piezoelectric transducer was affixed to the side of the PDMS using a two-component epoxy adhesive, ensuring the absence of gas bubbles within the glue and eliminating any air pockets between the piezo and the PDMS to prevent unwanted interference from gas. The transducer came equipped with two wires connecting its two faces, which were then linked to a function generator. Activation was carried out at a frequency of 240kHz, chosen as one of the resonant frequencies of the transducer. Voltage levels between 1VPP and 20VPP were applied during activation. The acoustic response was visualized using an optical microscope and recorded using high-sensitivity cameras.

### Numerical simulation

We conducted the numerical simulations by considering compressible Navier-Stokes equations and employing a perturbation expansion of the flow variables to obtain separate system of equations governing the system’s acoustic and mean response (*39*–*42*). The numerical results were obtained using the commercial finite element solver, COMSOL Multiphysics. The domain was discretised via triangular mesh elements, and we employed a mixed finite element scheme with Lagrange polynomials for the pressure and the velocity fields of order 1 and 2, respectively.

We consider a single spherical microbubble of 10 µm diameter, encapsulated in a 20 µm diameter oil droplet. The bubble-oil droplet is surrounded by water. The gas-oil interface is acoustically actuated in a combined volumetric and translational motion with frequency 240 kHz. The interface between the oil and the surrounding fluid (water) is prescribed the continuity of stress and continuity of the Lagrangian fluid velocity. For the surrounding water, the shear viscosity is set at 1.002 mPa·s, the bulk viscosity at 3.09 mPa·s, the density at 998.2 kg/m^3^, and the speed of sound at 1482 m/s. For the oil, we consider the following parameters: viscosity=10 mPa·s, density=920 kg/m^3^, and the speed of sound =1126 m/s.

## Acknowledgments

The authors thank

## Author contributions

Conceptualization: DA, NM, ADCF

Methodology: ADCF, DB, KGK

Investigation: ADCF, DB, NI, KGK

Supervision: DA, NM

Writing—original draft: ADCF, DA, NM

Writing—review & editing: ADCF, DA, NM

## Data and materials availability

All data are available in the main text or the supplementary materials.

