## Supplementary Information for "Programmable Assembly and Steering of Microbubble Droplets using Ultrasound"

Alexia Del Campo Fonseca *et al.*

**This PDF file includes:**

Figs. S1 to S8
Movies S1 to S4

**Other Supplementary Materials for this manuscript include the following:**
Movies S1 to S4

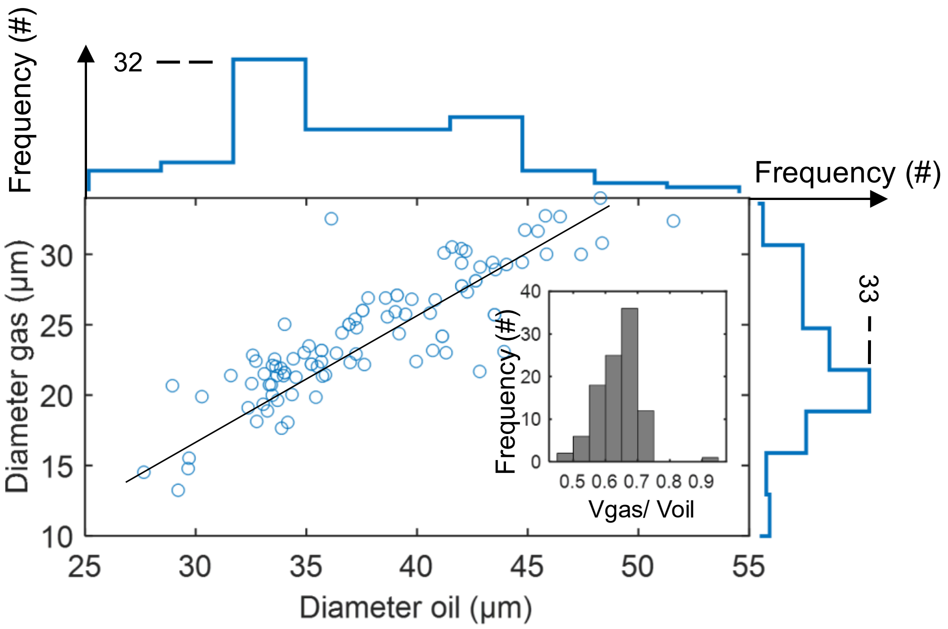


**Fig. S1.**

Size distribution of droplet composition is illustrated. The main plot displays the relationship between the diameter of the gas within ultrasound-responsive droplets and the diameter of the corresponding oil droplets. The side plots present the size histograms for both oil and gas, where the y-axis indicates the number (#) of droplets counted for each size. Additionally, the inset plots show a histogram depicting the frequency of each gas-to-oil ratio for droplets that remained stable and responsive to ultrasound.

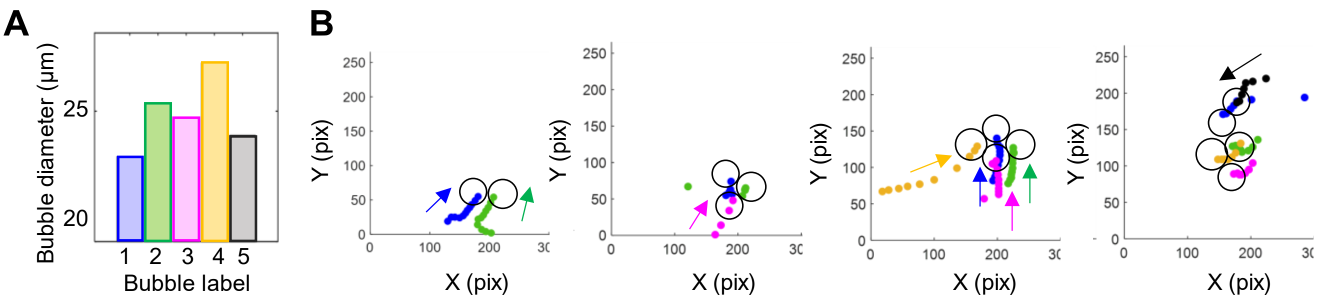


**Fig. S2.**

Analysis of a group of 5 microbubble droplets during self-assembly. This data corresponds with the results shown in Fig. 2 of the main manuscript. **(A)** Representation of the size of each bubble present in our target system. Colors correspond to the colors in Fig. 2, where blue is number 1, green is 2, pink is 3, yellow is 4 and black is 5. **(B)** These plots show the tracking of the motion of the bubbles over4 consecutive  time frames, on their way towards self-assembly.

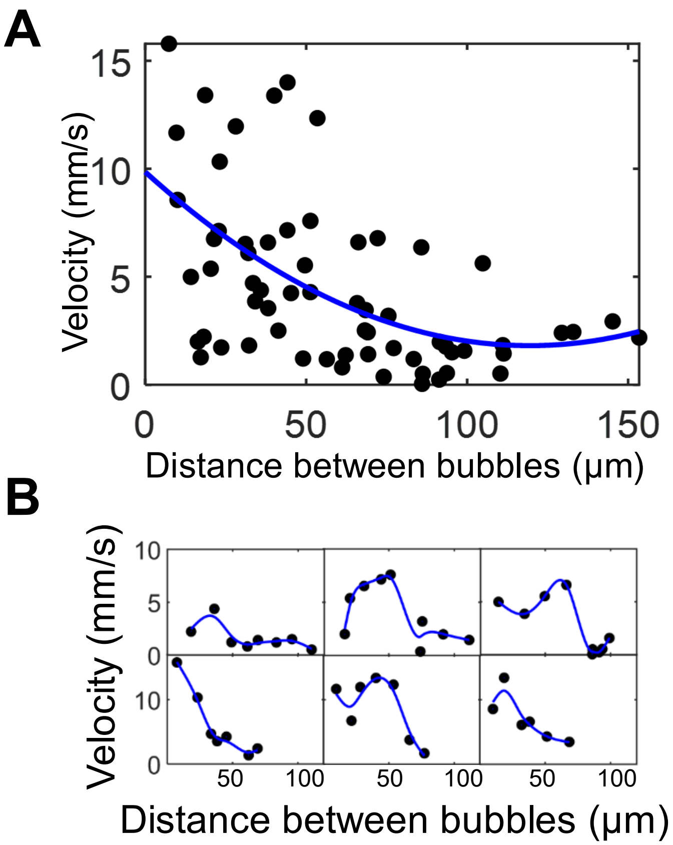


**Fig. S3.**
Measurement of droplet velocities during self-assembly experiments. **(A)** The velocity of droplets is plotted against the distance between neighboring bubbles. Due to significant variation in the distance to the transducers and the neighboring droplets, a wide range of responses is observed. To address this, individual droplet velocities are presented in the plots in B. **(B)** consists of an array of six plots, each representing a different droplet in separate experiments, all showing their velocities while they moved towards neighboring droplets.

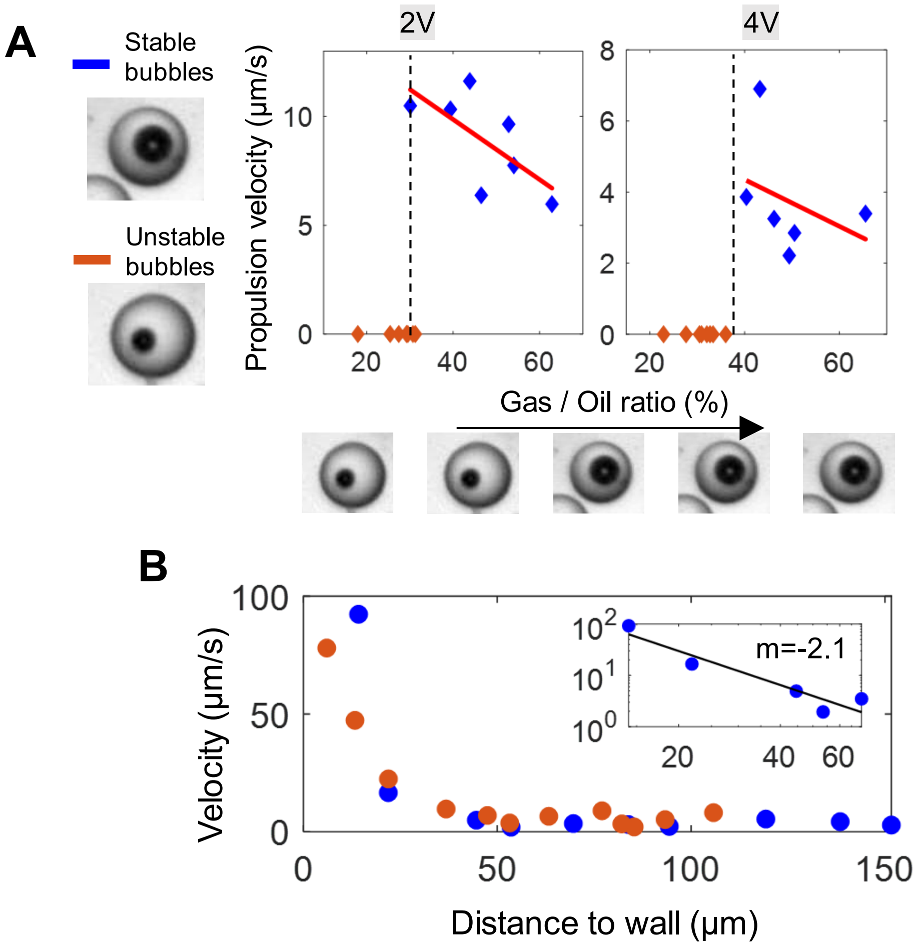


**Fig. S4.**

Analysis of droplet behavior during acoustic activation and in the presence of boundaries. **(A)** These plots illustrate the necessary droplet and bubble sizes required to maintain stable microbubbles during ultrasound activation. A low gas-to-oil ratio results in no droplet movement, leading to zero velocity and unstable bubbles. **(B)** This section represents the velocity of droplets relative to their distance from the wall, measured during the movement driven by the attraction between droplets and the channel wall, influenced by secondary radiation forces. These forces are proportionate to a factor of 2, as demonstrated in the inset plot, which provides a logarithmic analysis of the data points from the main plot.

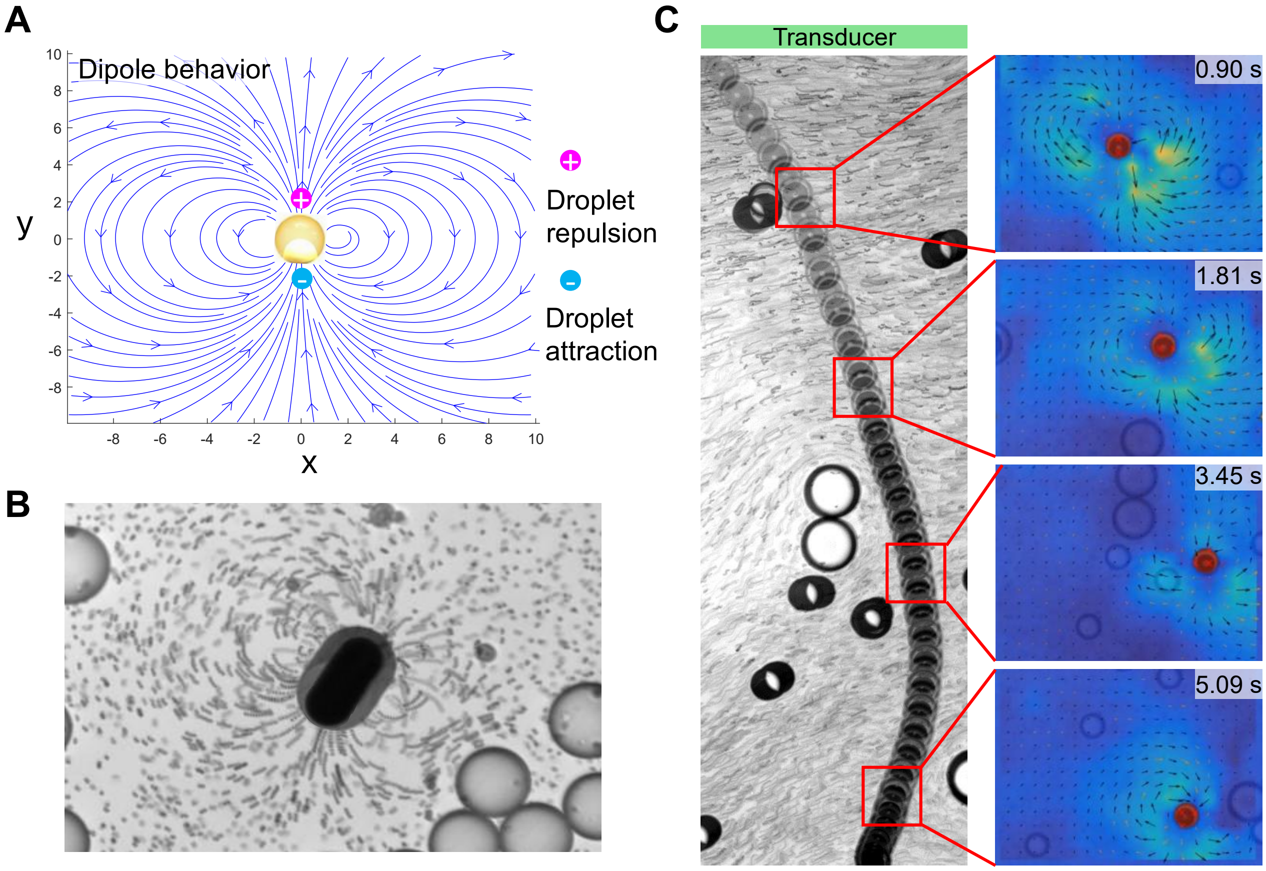


**Fig. S5.**

Analysis of acoustic streaming generated by microbubble droplets. **(A)** Visual depiction of the dipole-shaped streaming pattern produced during acoustic activation. **(B)** Experimental results captured with an optical microscope, showing a time-lapsed superposition of images (corresponding to the same image as in Fig. 4 of the main manuscript**). (C)** Further explanation of Fig. 4d: In Fig. 4d, each image represents the microbubble droplet at different distances from the transducer. This image visually illustrates these distances by showing the droplet's motion over time. The image on the left is also a superposition of images over a time frame of 6 seconds.

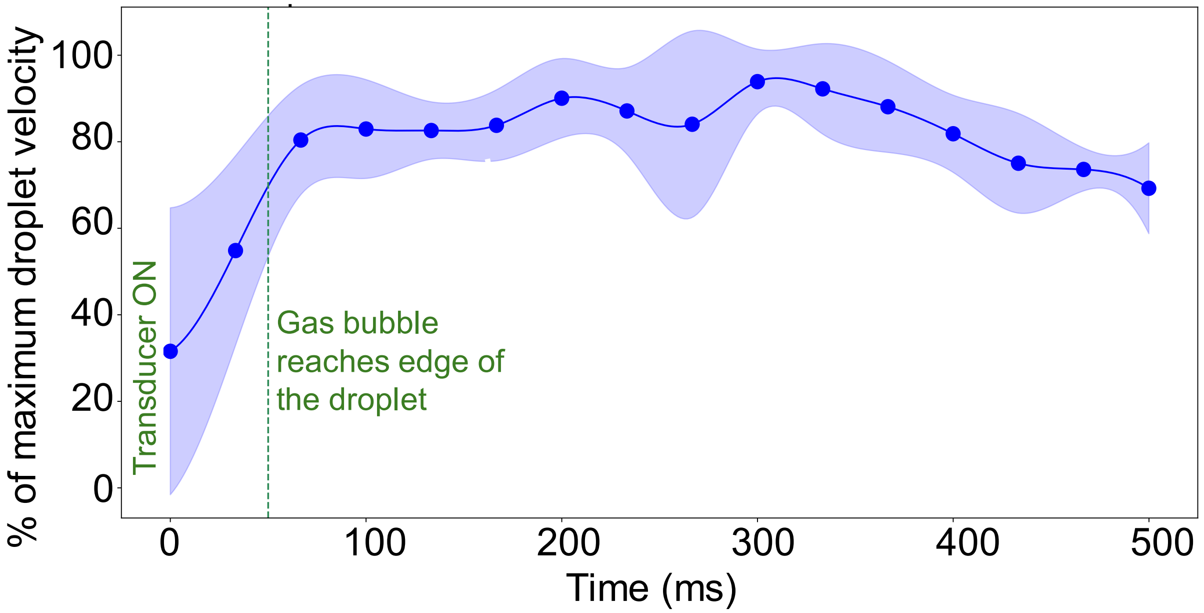


**Fig. S6.**

Analysis of acoustic streaming for microbubble droplet propulsion. This plot shows the normalized velocity of droplets in terms of % of maximum droplet velocity, over time. The shaded area represents the standard deviation at each time. The green shaded line marks the point where the gas bubble touched the edge of the droplet for each case.

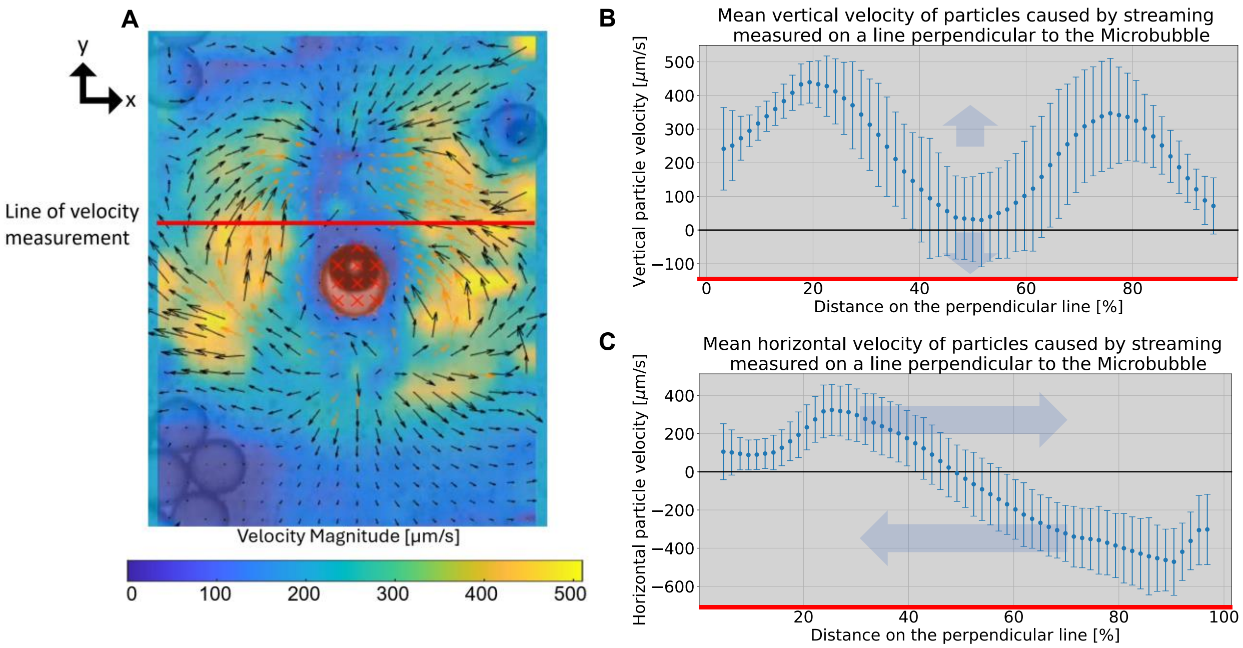


**Fig. S7.**

Analysis of streaming velocities. **(A)** Representation of PIV lab MATLAB analysis of the streaming generated around a microbubble droplet. The colors correspond to the colormap below and it represents the velocities of the streaming flow. **(B)** Vertical components of the streaming velocity following the length of the red line in ‘A’. **(C)** Horizontal components of the streaming velocity following the length of the red line in ‘A’.

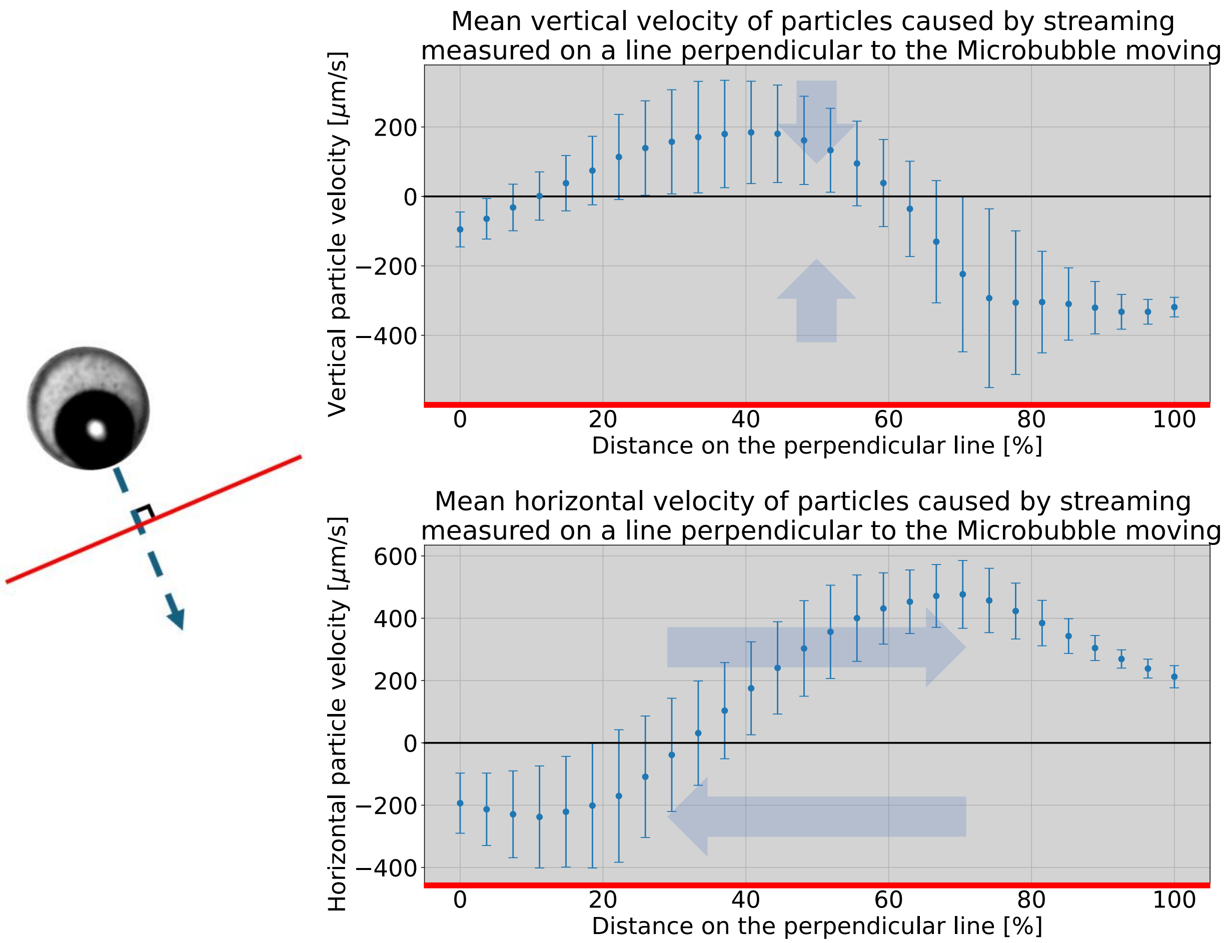


**Fig. S8.**

Analysis of streaming around a propelling droplet. The red line is positioned perpendicular to the direction of motion of the droplet. First plot shows the vertical components of the streaming velocity following the length of the red line. The second plot shows the horizontal components of the streaming velocity following the length of the red line.

**Movie S1.**

Droplets assembly into ordered structures. The assembly process of droplets is shown overt time under an acoustic field of 242 kHz and 20 $V_{PP}$.

**Movie S2.**

Gas bubble steering mechanism. In each video, the gas first aligns with the direction of motion, followed by the propulsion of the droplet in the desired direction. We have leveraged this for steering mechanism.

**Movie S3.**

Droplets fast self propulsion. This video shows on the left the self propulsion example of a single microbubble droplet under a sound field of 242 kHz and 20$V_{PP}$. On the background, we have placed 1µm tracer particles, in order to visualize the streaming flows generated. On the left, we have shown the PIV analysis of the moving tracer particles. This analysis correspond to the images and the colormap in Fig 4d.

**Movie S4.**

Propulsion and steering of microbubble droplets. By combining the activation of transducers at different locations, we first steer the inner gas bubble in the direction of motion, and consecutively we propel the droplet in the defined direction.
